# The Basolateral amygdala → Nucleus Accumbens core circuit mediates the conditioned reinforcing effects of cocaine-paired cues on cocaine seeking

**DOI:** 10.1101/2020.05.05.078329

**Authors:** Mickaël Puaud, Alejandro Higuera-Matas, Paul Brunault, Barry J. Everitt, David Belin

**Affiliations:** Department of Psychology, University of Cambridge, UK; Department of Psychobiology, School of Psychology. UNED. Madrid, Spain; CHRU de Tours, Équipe de Liaison et de Soins en Addictologie, Tours, France; UMR 1253, iBrain, Université de Tours, Inserm, Tours, France

**Keywords:** Basolateral amygdala, nucleus accumbens core, cocaine, second order schedule of reinforcement, conditioned reinforcement, chemogenetics

## Abstract

Individuals addicted to cocaine spend much of their time foraging for the drug. Pavlovian drug-associated conditioned stimuli exert a major influence on the initiation and maintenance of drug seeking often long into abstinence, especially when presented response-contingently, acting as conditioned reinforcers that bridge delays to drug use. The acquisition of cue-controlled cocaine seeking has been shown to depend on functional interactions between the basolateral amygdala (BLA) and the core of the nucleus accumbens (NAcC). However, the precise neuronal circuits underlying the acquisition of cue-controlled cocaine seeking behaviour have not been elucidated. Here we used a projection-specific Cre-dependent DREADD-mediated causal approach to test the hypothesis that the direct projections from the BLA to the NAcC are required for the acquisition of cue-controlled cocaine seeking behaviour. In Sprague Dawley rats with cre-mediated expression of the inhibitory DREADD Hm4Di in the NAcC projecting BLA neurons, treatment with CNO, but not vehicle, selectively prevented the impact of cocaine-associated conditioned reinforcement on cocaine seeking under a second-order schedule of reinforcement. This effect was attributable to the chemogenetic inhibition of the NAcC projecting BLA neurons as it was reversible, and absent in CNO-treated rats expressing an empty control virus. In contrast, chemogenetic inhibition of the anterior insula, which receives collateral projections from NAcC projecting BLA neurons, was without effect. These data demonstrate that the acquisition of cue-controlled cocaine seeking that depends on the conditioned reinforcing effects of cocaine cues require activity in the direct projections from the basolateral amygdala to the nucleus accumbens core.

## Introduction

Individuals with severe substance use disorder do not simply take drugs, they also spend a considerable amount of their time seeking and obtaining them. Over a prolonged history of drug use, this drug seeking becomes compulsive, persisting despite adverse personal as well as social consequences (1). It is therefore important to understand the neural basis of both drug seeking and drug taking behaviour, which are mediated by dissociable psychological processes (2–4).

Moreover, instrumental drug-seeking behaviour is greatly influenced by drug-associated Pavlovian conditioned stimuli (CS) (5). When presented unexpectedly and non-contingently, these drug cues capture attention (6), elicit approach behaviour (7, 8) and invigorate instrumental behaviour through the process of Pavlovian-Instrumental transfer (PIT; (9–11). However, it is when CSs response-produced, acting as conditioned reinforcers (CRfs), that they exert their most powerful effects on drug seeking behaviour, maintaining it over extended periods of time (12–15), precipitating relapse after abstinence (16–22) and increasing in impact the longer the period of abstinence (incubation of craving (23)).

At the neural systems level, the effects of self-administered cocaine that reinforce taking responses depend on activity in the mesolimbic dopamine system (3, 24), and especially dopaminergic transmission in the shell of the nucleus accumbens (NacS) (13). In marked contrast, the acquisition of cocaine seeking, in which responding maintained over protracted time periods is strongly enhanced by the presentation of cocaine-associated conditioned reinforcers (12–15, 25, 26), requires the functional integrity of the basolateral amygdala (BLA), the nucleus accumbens core (NAcC) and putative circuit interactions between these structures (27, 28).

Thus, studies using excitotoxic lesions or pharmacological manipulations have shown that the acquisition of cue-controlled cocaine seeking behaviour, as well as seeking responses for CRfs associated with food (29–31) and sex reward (32) depends on the basolateral amygdala (BLA) (33) which mediates the motivational representation of CS-US associations (9), and the NAcC, but not the NAcS (27). Functional disconnection studies further revealed that cue-controlled cocaine seeking behaviour depends on dopamine-dependent, interactions between the BLA and the NAcC (28), since unilateral blockade of dopamine receptors in the BLA combined with blockade of AMPA receptors in the contralateral NAcC, thereby functionally disconnecting these structures, impaired cue-controlled cocaine seeking behaviour to the same extent as bilateral manipulations of either structure alone (28).

While these studies suggest an important function of the BLA and the NAcC and their functional interaction in the acquisition of cue-controlled cocaine seeking, they do not precisely identify the circuit involved. However, glutamatergic BLA neurons exert robust physiological control over the activity of NAcC medium spiny neurons associated with reward seeking behaviour (34) and undergo synaptic plasticity following cocaine exposure (35), thereby suggesting that direct BLA→NAcC projections comprise the circuit that is required for the acquisition of cue-controlled cocaine seeking.

To test this hypothesis, we deployed a Cre-dependent, pathway-specific DREADD-mediated approach to causally interrogate the role of the BLA→NAcC circuit in the acquisition of cue-controlled cocaine seeking behaviour in Sprague Dawley rats.

The results show that the direct BLA→NacC circuit mediates the acquisition of cue-controlled cocaine seeking measured under a second-order schedule of reinforcement (SOR) (26). Thus, CNO administration prevented the ability of a cocaine-associated CRf to potentiate instrumental seeking responses in rats expressing the inhibitory hM4D(Gi) DREADD, but not an empty control virus, in the BLA→NacC neurons. This inhibition of cocaine seeking was reversible and was not observed after chemogenetic inhibition of the anterior insula to which the NAcC-projecting BLA neurons send collateral afferents (36).

## Materials and Methods

### Animals

Male Sprague Dawley rats (n=59, Charles River, Kent, UK) weighing approximately 300g upon arrival were housed in pairs for a week of habituation to the vivarium under a 12-hour reverse light/dark cycle (lights off at 07:00 am). Experiments were performed during the dark phase, 6-7 days per week, and were conducted in accordance with the United Kingdom 1986 Animals (Scientific Procedures) Act following ethical review by the University of Cambridge Animal Welfare and Ethical Review Body (AWERB) under the project licence number 70/8072.

### Procedures

A schematic of the timeline of the experiments is presented **Fig. 1A&C**. Briefly, after habituation to the animal facility rats underwent stereotaxic surgeries and following recovery they were singly housed. They were then implanted with an indwelling catheter in their right jugular vein (37) and, following recovery, were fed daily with 15-20g of standard chow per day prior to being trained to self-administer cocaine. Water was freely available in the home cage.

**Figure 1:**
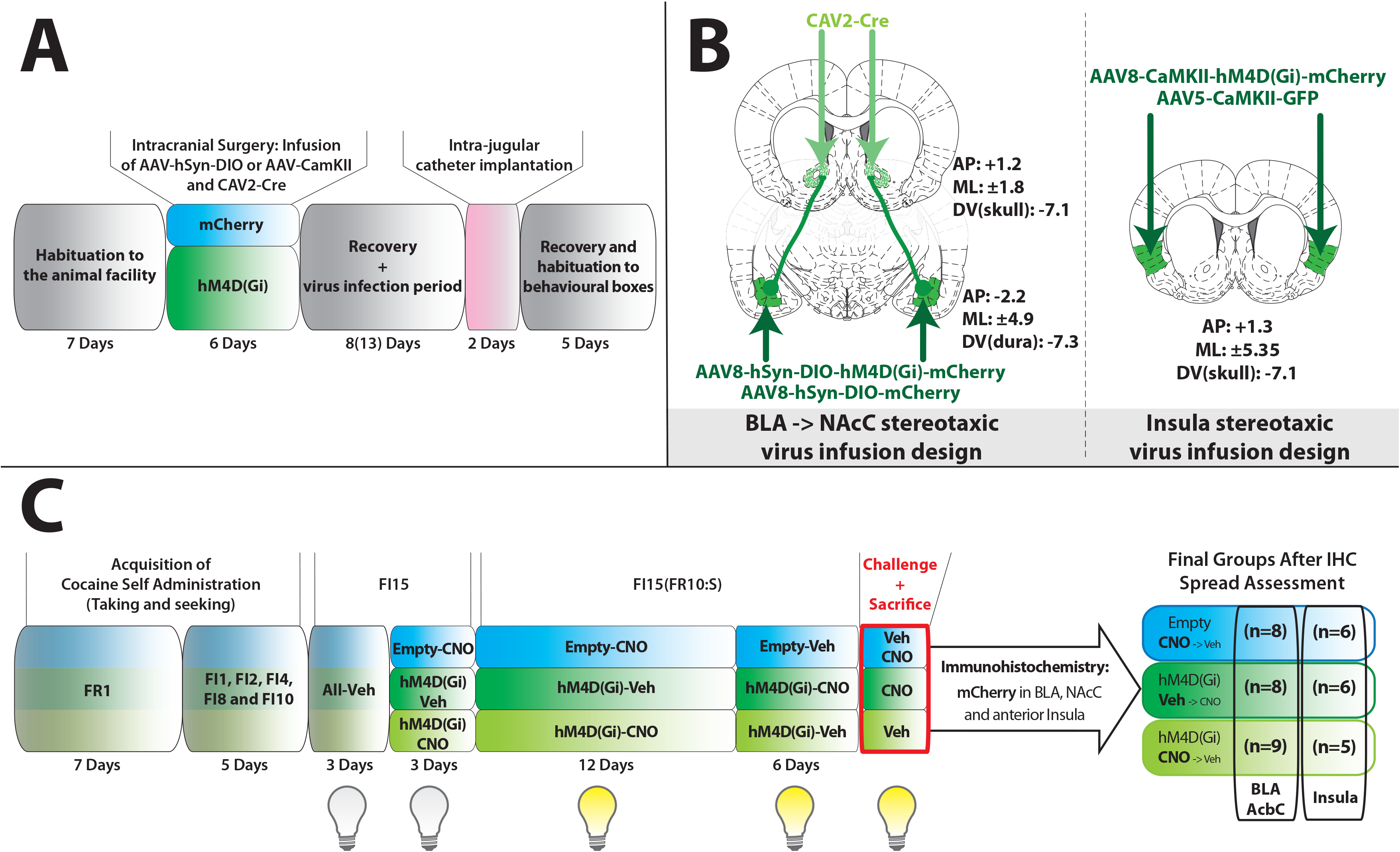
Timeline and experimental design. After a week of habituation to the vivarium (**A**) rats underwent intracranial surgeries during which they received virus infusions (**B**) enabling a projection-specific expression of empty control or Hm4D(Gi) DREADD in the NacC-projecting BLA neurons or in the anterior insula, to which NacC-projecting BLA neurons send massive collateral projections. After 8 to 13 days post-surgery rats were implanted with an indwelling catheter into their right jugular vein and allowed 5 days prior to the initiation of self-administration training (**C**). The various experimental groups, shown here in a colour-code used throughout, were initially trained to acquire cocaine self-administration, under continuous reinforcement (fixed ratio 1, FR1) for seven days. Rats were then progressively trained to respond under fixed interval schedules of reinforcement (FI), from 1 min to 10 min over 5 sessions. They were then trained to seek cocaine for three days under a FI15 schedule of reinforcement (FI15) prior to receiving either veh or CNO treatment daily, for three additional FI15 sessions and 12 days of responding under a FI15(FR10:S) second order schedule of reinforcement in which cocaine seeking responses are reinforced every tenth lever press by the contingent presentation of the cocaine-paired cue acting as a conditioned reinforcer. Treatment was subsequently reversed for six additional days of training under FI15(FR10:S) after which rats were deeply anaesthetized and perfused brains harvested for immunohistochemical assessments.

### Drugs

Cocaine hydrochloride (NIDA Drug Supply Programme) was dissolved in sterile 0.9% NaCl. Clozapine-N-oxide (NIDA Drug Supply Programme) was dissolved first in 5% DMSO (Dimethyl Sulfoxide, Sigma-Aldrich, Poole, UK) and then in sterile 0.9% NaCl (Henry Schein Ltd, UK). A vehicle solution was prepared with 5% DMSO in sterile 0.9% saline as a control for CNO.

### Viral vectors

Cre-dependent expression of hM4D(Gi) was mediated by a co-administration of a trans-synaptic CAV2-Cre virus (Plateforme de Vectorologie de Montpellier (PVM)) and a pAAV8-hSyn-DIO-hM4D(Gi)-mCherry virus (Addgene; plasmid #44362) while that of the mChrerry reporter alone was mediated by co-administration of the CAV2-Cre virus and a pAAV8-hSyn-DIO-mCherry virus (Addgene; plasmid #50459) called ‘Empty’ throughout. Non-Cre-dependent expression of hM4D(Gi) was mediated by administration of an AAV8-CaMKIIa-hM4D(Gi)-mCherry virus (Addgene; plasmid #50477) while that of a GFP reporter by administration of an AAV5-CaMKII-GFP virus (Boyden, UNC Vector Core) also called ‘Empty’ throughout.

Upon arrival, all viral vectors were aliquoted and stored at −80°C until subsequent use. All viruses were freshly diluted in sterile PBS at a final concentration of 1×10^9^vp/uL.

### Stereotaxic surgery and viral infusions

Viral infusions (design illustrated **Fig. 1B)**, were performed at least 29 days before the first chemogenetic manipulation thereby enabling the expression of the transgenes, which was assessed immediately after the last challenge.

The CAV2-Cre virus was infused bilaterally (10^9^vp/μl, 1μl/side) into the NAcC at the following stereotaxic coordinates (Paxinos and Watson 2009) AP: +1.2, ML: ±1.8, −7.1 DV (from skull) (38). The AAV8-hSyn-DIO-hM4D(Gi)-mCherry and AAV8-hSyn-DIO-mCherry viruses were infused bilaterally (1ul/side) into the BLA at the following stereotaxic coordinates AP: −2.5, ML: ±4.9, DV (from dura): −7.2 DV (from dura) (39). The AAV8-CaMKIIa-hM4D(Gi)-mCherry and AAV5-CaMKII-GFP viruses were infused bilaterally (0.85ul/side) into the anterior insular cortex at the following stereotaxic coordinates AP: +1.3, ML ±5.35, −7.1 DV (from skull) (40). All infusions were performed with 10ul Hamilton syringes placed in a Harvard infusion pump and connected with a polyethylene tubing to 24 gauge injectors (Coopers needle works Ltd) at a rate of 0.15 μl/min. Injectors were left in place for 7min following completion of the infusion to allow for diffusion away from the injector tip.

### Intra-jugular catheterisation Surgery

Rats were implanted with a home-made indwelling catheter into their right jugular vein under isoflurane anaesthesia (O_2_: 2L/min; 5% for induction and 2-3% for maintenance and analgesia (Metacam, 1mg/kg, sc., Boehringer Ingelheim) as previously described (41). Following the surgery, rats received daily oral treatment with the analgesic for three days and an antibiotic (Baytril, 10mg/kg, Bayer) for a week. Catheters were flushed with 0.1 ml of heparinized saline (50 U/ml, Wockhardt®) in sterile 0.9% NaCl every other day after surgery and then before and after each daily self-administration session.

### Apparatus

Experiments were conducted in 24 standard operant chambers (Med Associates Inc., St. Albans, VT, USA) controlled by MedPC software (Med Associates Inc., Ltd) as previously described (42). Chiefly each (29.5 × 32.5 ×23.5 cm) chamber was housed in ventilated, sound-attenuating cubicles. Sidewalls were aluminium; the ceiling, front and back walls were clear polycarbonate. Two retractable levers (4cm wide) were situated 8cm above the grid floor and 12cm apart; a white cue light (2.5W, 24V) was situated above the levers and a white house light (2.5W, 24V), situated on the wall opposite the levers. Implanted catheters were connected to a 10mL syringe driven by an infusion pump (Semat Technical, Herts, UK) via Tygon tubing itself protected within a spring leash attached to a swivel connected to a balanced metal arm secured outside of the chamber.

### Self-administration

Rats were trained to self-administer cocaine (0.25□mg/100μl/5.7s/infusion) under continuous reinforcement (Fixed Ratio 1 (FR1)) over 7 daily 2-hour sessions. Under this schedule, each active lever press resulted in drug infusion initiated concurrently with a 20s time out that included onset of a 20s illumination of cue light positioned above the active lever (conditioned stimulus; CS), offset of the house light and retraction of both levers. Inactive lever pressing was recorded but had no scheduled consequence. Active and inactive lever assignment was counterbalanced, and a maximum of 30 infusions was available for this stage.

Following these 7 daily sessions under continuous reinforcement, the daily schedule of reinforcement was changed to fixed intervals, increasing across daily training sessions from 1 min (fixed interval 1 min, FI1) to FI2, FI4, FI8, FI10 and eventually FI15 min (38). After three sessions on FI15 before which rats were habituated to daily IP injection of vehicle (1ml/kg), rats were tested for their drug seeking behaviour under FI15 over three daily sessions following IP administration of either vehicle or CNO (5mg/kg i.p.).

Having tested the influence of chemogenetic inhibition of the BLA→NacC pathway on responding for cocaine under FI15, the role of the pathway on the impact of conditioned reinforcement of the cocaine-associated CS was measured over 12 daily sessions. Thus, each tenth lever press resulted in the contingent presentation of the CS for 1 second, while cocaine (and the associated CS) was delivered on the first tenth lever press after a 15 min interval had elapsed; formally this is a second-order schedule of reinforcement (SOR) of the type FI15(FR10:S). Rats were trained to respond over 2-hour sessions or for five cocaine infusions. CNO or vehicle were administered prior to each daily session as described above, and, after 12 SOR sessions, the reversibility of the CNO-induced inhibition of the BLA→NacC pathway in hM4D(Gi)-expressing rats was tested over 6 additional sessions during which those hM4D(Gi)-expressing that had previously received CNO received vehicle and vice-versa.

### Tissue collection

Ninety minutes following a 15min drug-free seeking session under CNO vs Veh treatment rats were deeply anaesthetised with pentobarbital (Euthatal, Merial, 750mg/Kg) and perfused with 0.01M phosphate-buffered saline (PBS), followed by 4% neutral-buffered formaldehyde (NBF). Brains were collected and post-fixed for at least 24h at 4°C in 4% NBF. Brains were then cryo-protected in a 30% sucrose solution (prepared in PBS 0.01M). After quick freezing on dry-ice, brains were processed into 35μm coronal sections using a cryostat (Leica Microsystems). Sections were kept free floating in a cryoprotectant solution (30% sucrose, 30% ethylene glycol, 0.547% Na_2_HPO_4_, 0.159% NaH_2_PO_4_, 0.9% NaCl, 1% polyvinyl pyrolidone, in distilled water) and stored at −20°C until immunohistochemistry.

### Immunohistochemistry

Brain sections were washed three times 10min in 0.01☐M PBS at room temperature. Sections were then blocked for 2☐h in 5% bovine serum albumin (BSA, Sigma-Aldrich, A7906) in 0.01☐M PBS and 0.3% Triton X-100 (Sigma-Aldrich, T8787) prior to being incubated with the primary antibody (rabbit anti-mCherry; 1:1000; abcam, ab167453) in a 2% BSA and 0.1% Triton X-100 overnight (18h) at 4°C. Sections were then washed three times for 10min each in 0.01☐M PBS and incubated in secondary antibody (goat Alexa Anti-Rabbit 488, 1:1,000; ThermoFisher Scientific, #A-11008) for 2☐h at room temperature. Sections were again washed three times 10min with 0.01☐M PBS and mounted onto glass slides (Fisherbrand Superfrost Microscope Slides) and allowed to dry overnight (protected from light). Slides were then covered with a coverslip and fluoroshield mounting medium (abcam, ab104135). Slides were stored at 4°C prior to image acquisition. Images were acquired with a Zeiss Axio Imager M2 equipped with an AxioCam MRm camera (Oberkochen, Germany). Images were taken using Visiopharm® software (Medicon Valley, Denmark), at magnification 5x and tiled to create the whole slices images or at magnification 10x for the regions of interest.

### Data and statistical analyses

Data, analysed using STATISCA 10 (Statsoft, Palo Alto, USA), are presented as means ± SEM or box plots [medians ±25% (percentiles) and Min/Max as whiskers)].

Analyses were performed on 41 rats as 10 rats were excluded following histological assessment of the spread of the expression of the reporters and 8 rats were lost from the study because of catheter failures.

Assumptions for normal distribution, homogeneity of variance and sphericity were verified using the Shapiro–Wilk, Levene, and Mauchly sphericity tests, respectively. When assumptions were violated data were log-transformed. However, for the sake of transparency, both non transformed and transformed data are presented throughout.

Lever presses during acquisition of both cocaine self-administration and cocaine seeking across the increasing duration of the Fixed Interval schedules of reinforcement, were analysed using a repeated-measures analysis of variance (ANOVAs) with lever (active and inactive) and sessions as within☐subject factors (**Fig. S1 and S2**), and treatment (Hm4D(Gi)-CNO, Hm4D(Gi)-Veh and Empty-CNO) as between-subject factor.

Because of between-session variability in performance across groups, data pertaining to the acquisition of cue-controlled cocaine seeking were analysed across 5 3-day blocks (1 FI 15 block and 4 SOR blocks) using 2-Way ANOVAs with blocks as within-subject factors and treatment as between-subject factors.

The data shown as boxplots were analysed using a one-way ANOVA with blocks as within-subject factors.

Significant interactions were analysed further using Duncan’s *post hoc* analyses or hypothesis-driven planned comparisons (DREADD-CNO should prevent potentiation of responding by CRf as compared to the two control groups together) wherever appropriate.

For all analyses, significance was set at α = 0.05. Effect sizes are reported as partial eta squared (p*η*^2)^

## Results

The expression of inhibitory Hm4D(Gi) DREADDs or empty-virus controls, the reporters of which were heavily expressed in cell bodies and axon terminals of NAcC-projecting BLA neurons (**Fig. 2A**), had no effect on the acquisition of cocaine self-administration under continuous reinforcement (fixed ratio 1, FR1) (**Fig S1A**) or on cocaine seeking under fixed interval schedules of reinforcement in the absence of presentation of cocaine-associated CRfs. Thus, the three experimental groups showed similar increase in their active lever presses (AL) over the seven daily sessions under FR1 [main effect of session: F_6,132_=2.25, p=0.042, p*η*^2^=0.09 and group x session interaction: F_12,132_<1, p*η*^2^=0.06] and the subsequent 8 sessions under FI of increasing durations, from 1 min to 15 min [main effect of session: F_7,154_=28.94, p<0.01, p*η*^2^=0.57 and group x session interaction: F_14,154_=1.75, p=0.051, p*η*^2^=0.14] to that previously described (38, 39, 43), even if the empty controls tended to respond at a slightly higher rate than the two Hm4D(Gi) groups under FI.

**Figure 2:**
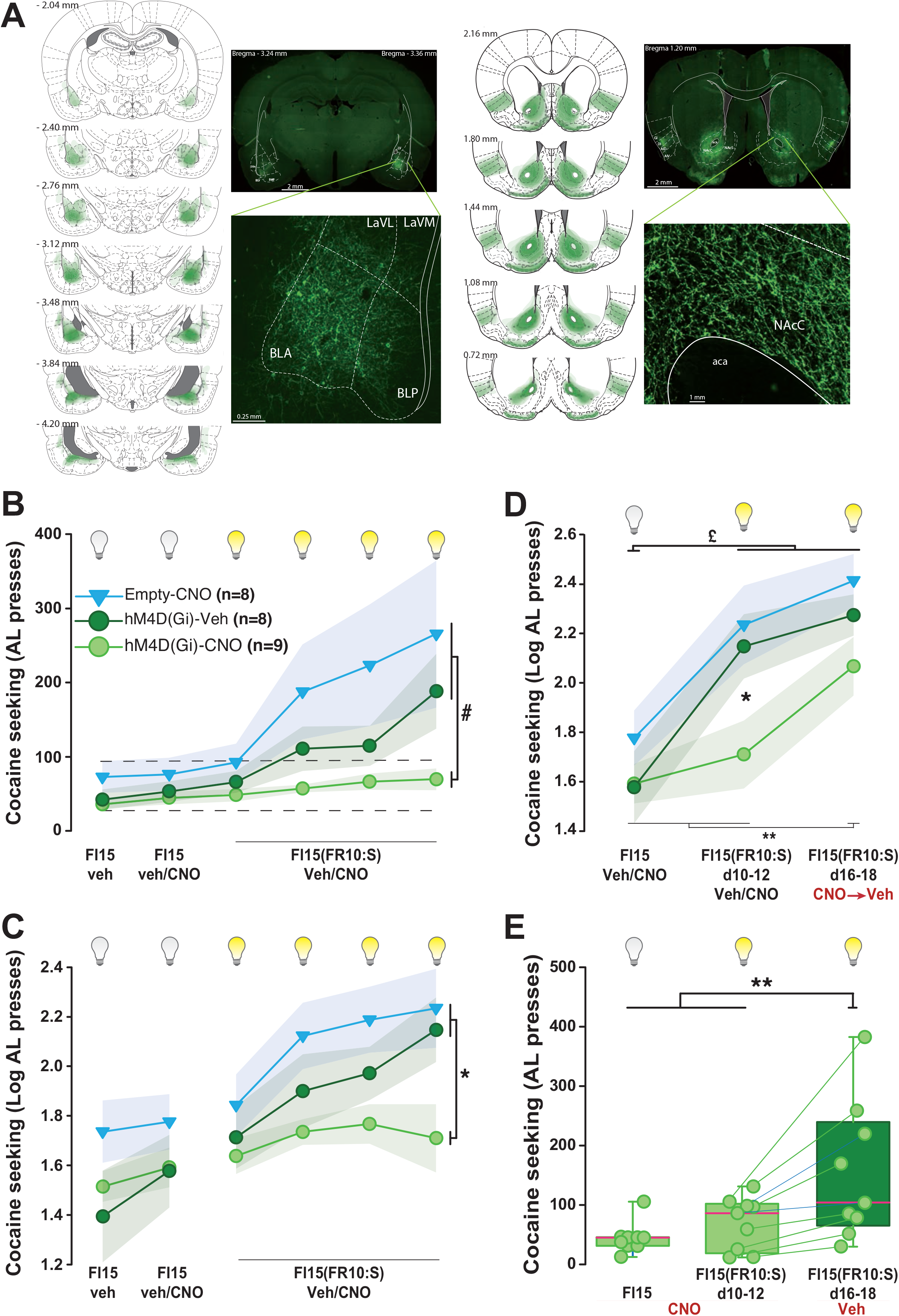
Chemogenetic inhibition of NAcC-projecting BLA neurons specifically prevents the potentiation of cocaine seeking behaviour by drug paired cues otherwise acting as conditioned reinforcers. **A)** Cre-mediated projection-specific expression of Hm4d(Gi) in the NacC-projecting BLA neurons resulted a dense expression of the reporter both in the cell bodies and terminals of targeted neurons as shown in representative photos of hM4D(Gi) expressing neurons (mCherry tag revealed by immunofluorescence in green) in the BLA (left) and their axon terminals in the NAcC (right). Alongside is a schematic of coronal sections of the brain covering the anterior-posterior extent of the BLA (left) or the NAcC (right) with density maps depicting in green the spread of mCherry expression in each structure across all individuals. (LaVL: lateral amygdala ventrolateral, LaVM: lateral amygdala ventromedial, BLA: basolateral amygdala anterior, BLP: basolateral amygdala posterior, NAcC: Core of the nucleus accumbens, aca: anterior commissure). The majority of rats displayed a labelling restricted to the NacC, with strong labelling in the anterior insula to which NAcC-projecting BLA neurons send collateral projections. Some rats had expression spreading to the dorsolateral part of the nucleus accumbens shell and the piriform cortex. **B-C)** The introduction of the cocaine-paired cue contingently upon responding resulted in a substantial increase in cocaine seeking over time in both the empty-CNO and Hm4D(Gi)-Veh groups during the first 15min drug-free period of daily sessions represented either as non- (**B**) or Log-transformed data (**C**). In marked contrast, such potentiation or responding was not observed in Hm4D(Gi)-CNO rats whose seeking responses were much lower than that of the two control groups over the four 3-session blocks of training under FI15(FR10:S) (# planned comparison: p<0.03) and actually never differed from that observed under FI15 (the dotted lines represented on **B** represent the SEM of the entire population during the first FI15 block). The effect of chemogenetic inhibition of the NacC-projecting BLA neurons was specific of the potentiation of cocaine seeking responses by the conditioned reinforcing properties of the CS as it had no effect on performance under FI15, prior to the introduction of the contingent presentations of the CS. The prevention of acquisition of cue-controlled cocaine seeking by chemogenetic inhibition of the NacC-projecting BLA neurons was reversible (**D-E**). Thus, upon reversal of treatment, e.g., when Hm4D(Gi)-CNO rats, whose drug seeking under FI15(FR10:S) was hitherto much lower than that of both control groups (*: post-hoc, p<0.05) and similar to that expressed under FI15, received vehicle in place of CNO, they immediately showed sensitivity to the conditioned reinforcing properties of the CS as they displayed a potentiation of responding which rate became, within a week, higher than that under FI15 (**, post-hoc: p<0.01) and indeed similar to that of the empty and Hm4d(Gi) control groups (**D**). Further analysis of the behaviour displayed by each individual of the Hm4d(Gi)-CNO→Hm4D(Gi)-Veh group revealed that the increase in responding shown at the group level upon reversal of treatment (as compared both to FI15 baseline and the last 3-day block of FI15(FR10:S) reflected an increase in lever press in each of the 9 individuals (**E**). Empty-CNO, hM4D(Gi)-Veh and Hm4D(Gi)-CNO are represented in blue triangles, dark green and light green circles, respectively.

The CNO-induced activation of inhibitory Hm4D(Gi) DREADDs had no effect on seeking responses under a FI15 schedule of reinforcement (**Fig. 2B & C**). Thus, over the first drug-free interval of three days of training under FI15 responding under the different treatments did not differ between groups (e.g. Hm4D(Gi)-Veh, Hm4D(Gi)-CNO, empty-CNO) or from baseline performance measured over three days prior to the introduction of treatment (**Fig 2C**) [main effect of block: F1,22=10.3, p=0.004, p*η*^2^=0.32, group: F_2,22_=1.35, p=0.28, p*η*^2^=0.11 and block x group interaction: F_2,22_=1.83, p=0.18, p*η*^2^=0.14].

However, as predicted, chemogenetic inhibition of NAcC-projecting BLA neurons prevented the potentiation of instrumental drug seeking responses, observed in empty-CNO and Hm4D-Gi-Veh control groups, following the introduction of contingent presentations of the cocaine-paired CS (**Fig 2B and C**) [main effect of block: F_4,88_=23.13, p<0.01, p*η*^2^=0.51 and group x block interaction: F_8,88_=2.67, p=0.011, p*η*^2^=0.19]. Indeed, Hm4D(Gi)-CNO rats failed to show an increase in active lever presses on introduction of the FI15(FR10:S) SOR and maintained a low level of responding similar to that seen under FI15 baseline conditions throughout. Consequently Hm4D(GI)-CNO rats significantly differed from the two control groups [planned comparison: F_1,22_=5.49, p<0.03] in which cocaine seeking steadily increased over the 4 3-day session blocks of training under FI15(FR10:S) eventually to reach levels twice as high as those displayed under baseline conditions (p<0.01).

As soon as the treatment was reversed and Hm4D(Gi)-CNO rats started to receive vehicle instead of CNO (Hm4D(Gi)-CNO→Veh), they quickly increased their cocaine seeking responses under the same FI15(FR10:S) conditions and they eventually reached levels of responding that were similar to that of the empty-CNO→veh and Hm4D(Gi)-Veh→CNO control groups, which maintained steady levels of cue-controlled cocaine seeking [main effect of group x block interaction: F_4,44_=3.04, p=0.027, p*η*^2^=0.22] (**Fig. 2D**). Thus, Hm4D(Gi)-CNO→Veh rats eventually showed the potentiation following introduction of response-contingent presentations of the cocaine-paired CS as compared to their previous performance under FI15 baseline (39, 43) and FI15(FR10:S) under CNO (post-hoc here) (**Fig. 2D**). The reversibility of the effect of the chemogenetic inhibition of the NAcC-projecting BLA neurons on the acquisition of cue-controlled cocaine seeking was further supported by an analysis of individual performance upon reversal of treatment (**Fig. 2E**). Thus, each of the 9 rats in that group substantially increased their seeking behaviour for cocaine on reversal of CNO treatment [F_2,16_=9.33, p=0.002, p*η*^2^=0.54].

The introduction of CNO treatment in the Hm4D(Gi)-veh (Hm4D(Gi)-veh→CNO) group after a period of 18 days of training to seek cocaine had no effect on well-established cue-controlled cocaine seeking (**Fig. 2D**).

Taken together, these results show that the NAcC-projecting BLA neurons are necessary for the potentiation of cocaine seeking by the conditioned reinforcing properties of drug-paired cues. However, histological assessment of the pattern and spread of expression of the transgenes revealed that in addition to the dense expression in the BLA→NacC circuit, robust expression was systematically also observed in the anterior insular cortex (AI). This demonstrates that NAcC-pojecting BLA neurons send substantial collateral projections to the AI, confirming earlier observations already, but the functional significance of which was not investigated (36). We therefore investigated the possible involvement of the AI in Pavlovian mechanisms and instrumental conditioning (44, 45) by studying the influence of chemogenetic inhibition of NAc-projecting BLA neurons on the impact of conditioned reinforcement on cocaine seeking.

An independent cohort rats had virus-mediated expression of Hm4D(Gi) or reporter-only under the CAMKII promoter bilaterally in the AI (**Fig 1 and Fig 3A**) prior to being tested in the same procedure as that described above. The expression of inhibitory Hm4D(Gi) DREADDs or empty-virus controls, the reporters of which were heavily expressed in cell bodies of the AI (**Fig. 3A**), had no effect on the acquisition of cocaine self-administration under FR1 (**Fig S2A**) or cocaine seeking under fixed interval schedules of reinforcement. Thus, the three experimental groups showed similar increase in their active lever presses (AL) over the seven daily sessions under FR1 [main effect session: F_6,84_=8,32, p<0.01, p*η*^2^=0.37 and group x session interaction: F_12,84_<1, p*η*^2^=0.07) and the subsequent 8 sessions under FI of increasing durations, from 1 min to 15 min [main effect session: F_7,98_=22.95, p<0.01, p*η*^2^=0.62 and group x session interaction: F_14,98_<1, p*η*^2^=0.06] (**Fig S2B**) to that of those of the first experiment.

**Figure 3:**
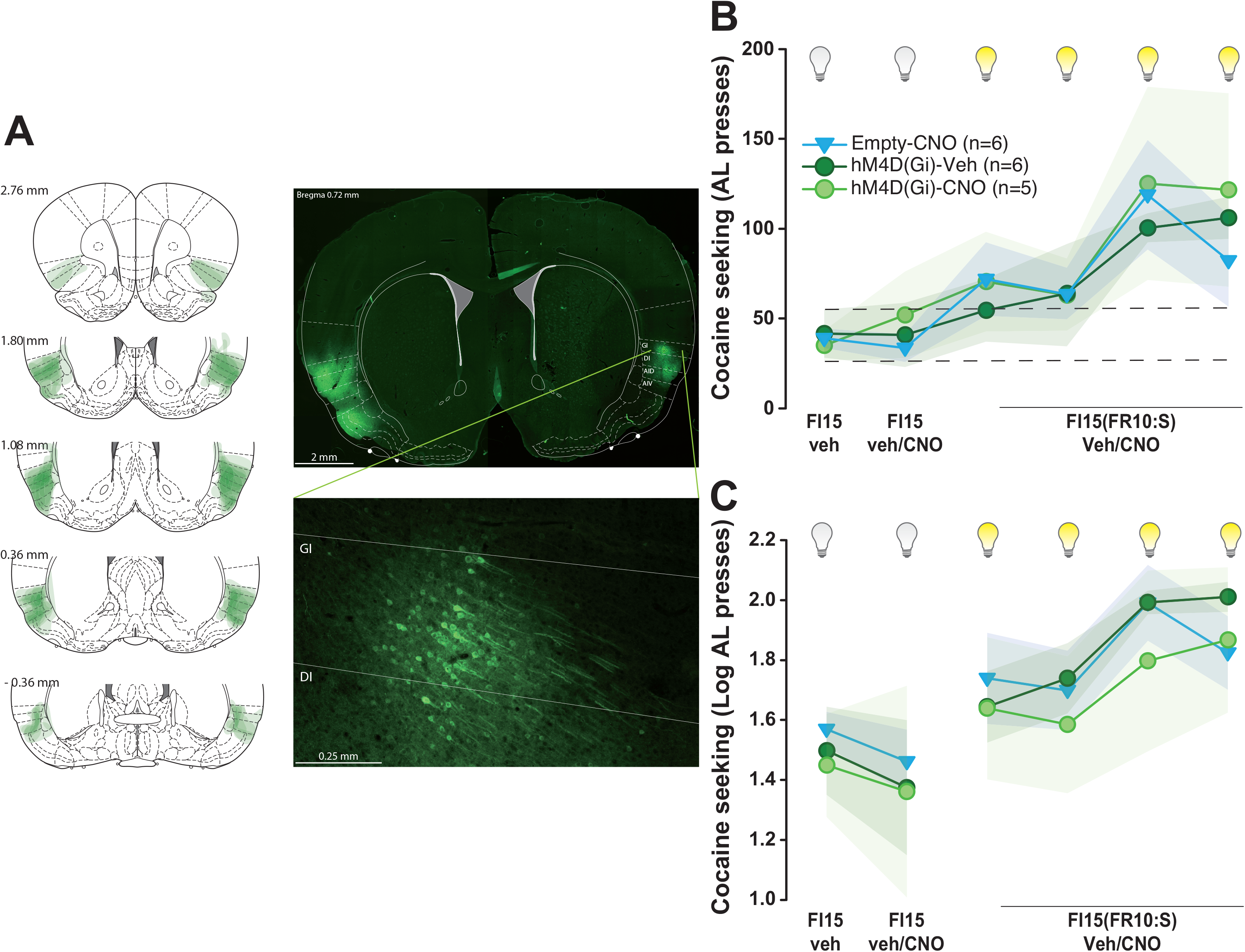
Chemogenetic inhibition of the anterior Insular cortex does influence the potentiation of drug seeking by cocaine-paired conditioned stimuli acting as conditioned reinforcers. **A)** Viral-mediated expression of Hm4d(Gi) in the anterior insula (AI) resulted a dense expression of the reporter in the cell bodies of targeted neurons as shown in representative photos of hM4D(Gi) expressing neurons (mCherry tag revealed by immunofluorescence in green) in the AI (left). Alongside is a schematic of coronal sections of the brain covering the anterior-posterior extent of the AI with density maps depicting in green the spread of mCherry expression in each structure across all individuals. A dense expression of the reporters was almost exclusively restricted to the AI, with some spread observed in some rats in adjacent territories in the dorsal part of the 3 layers of the piriform cortex, the dorsal endopiriform nucleus and the claustrum. **B-C**) Chemogenetic inhibition of the AI had no effect on cocaine seeking under FI15 or on the potentiation of drug seeking responses by the conditioned reinforcing properties of drug-paired CSs. Indeed, non- (**B**) or log-transformed (**C**) active lever presses measured during the first 15 min drug-free interval of 12 daily sessions represented in blocks of 3 sessions never differed between the Hm4D(Gi)-CNO and the two control groups, namely empty-CNO and Hm4D(Gi)-Veh. Thus, upon introduction of the FI15(FR10:S) second order schedule of reinforcement, all groups displayed a similar steady increase in responding as compared to FI15 baseline, thereby demonstrating their sensitivity to the conditioned reinforcing properties of the cocaine-paired CSs The dotted lines on B represent the SEM of the entire population for the first FI15 block. Empty-CNO, hM4D(Gi)-Veh and Hm4D(Gi)-CNO are represented in blue triangles, dark green and light green circles, respectively

As previously, the CNO-induced activation of inhibitory Hm4D(Gi) DREADDs had no effect on cocaine seeking under a FI15 schedule of reinforcement (**Fig. 3B-C**). Thus, cocaine seeking responses made during the first drug-free intervals over three days of training under FI15 did not differ between groups (e.g. Hm4D(Gi)-Veh, Hm4D(Gi)-CNO, empty-CNO) or from baseline performance measured over three days prior to the introduction of treatment (**Fig 3B-C**) [main effect of group: F_2,14_<1, p*η*^2^=0.01 and block x group interaction: F_2,14_<1, p*η*^2^=0.12].

However, in marked contrast with the effect of chemogenetic inhibition of NAcC-BLA projecting neurons, inhibition of the AI had no effect on the potentiation of cocaine seeking that accompanies response-contingent presentation of the drug-paired under a FI15(FR10:S) second order schedule of reinforcement (**Fig 3B-C**). Thus, on the introduction of the conditioned reinforcer, Hm4D(Gi)-CNO rats displayed the same increase in drug seeking responses as that shown by the Hm4D(Gi)-Veh and empty-CNO control groups [main effect of block: F_4,56_=11.82, p<0.01, p*η*^2^=0.46 and group x block interaction: F_8,56_<1, p*η*^2^=0.06].

## Discussion

Taken together, the data presented here reveal that the direct BLA→NAcC circuit mediates the impact of conditioned reinforcement on cocaine seeking. Thus, chemogenetic inhibition of the NAcC-projecting BLA neurons prevented the potentiation of responding that follows seeking response-contingent presentation of cocaine-associated CSs (26). Importantly, these effects are specific to the BLA→NAcC as chemogenetic inhibition of the anterior insula, to which NAcC-projecting BLA neurons send substantial collateral projections, had no effect on either instrumental seeking responses or their potentiation by cocaine cues.

These results extend understanding of the amygdalo-striatal mechanisms involved in conditioned reinforcement (13, 25, 46, 47), and particularly its impact on the seeking of stimulant drugs (5, 25, 47, 48).

Thus, drawing on early evidence that the BLA and its functional interactions with the NacC mediates the impact of conditioned reinforcers on instrumental responding for natural reinforcers such as water or a sexual partner (32, 49), studies using a variety of pharmacological and physical lesion-based manipulations of the amygdalo-striatal system, including functional disconnections, have provided evidence of a causal role of the BLA (31, 33), the NAcC (27) and their functional interaction in mediating the conditioned reinforcing properties of drug-paired CSs and their impact on the reinstatement of extinguished drug seeking instrumental responses (27, 28, 33, 50, 51). Bilateral BLA (33) or NAcC (but not NacS) (27) excitotoxic lesions prevented the acquisition of cocaine seeking under a second order schedule of reinforcement. Additionally functional disconnection of the BLA and the NAcC showed that coordinated dopaminergic activity in the BLA and glutamatergic activity in the NAcC is involved in the acquisition of cue-controlled cocaine seeking (28).

Here we extend these findings by showing that these functional interactions depend on a specific BLA→NAcC pathway whereby glutamatergic inputs from the BLA influence downstream processes in the NAcC to mediate the effects of the conditioned reinforcing properties of cocaine-paired cues on instrumental drug seeking behaviour.

This observation is consistent with the previous demonstration that BLA neurons gate, in a glutamate-dependent manner, the activity of NAcC medium spiny neurons and consequent reward-seeking behaviour (34), and that plasticity at the BLA→NAc synapse is involved in the acquisition of responding reinforced by the contingent presentation of a cocaine-paired cue after withdrawal (51).

In addition, the present results reveal that, unlike zif-268 knockdown-mediated long-lasting disruption of the reconsolidation of the CS-cocaine memory in the BLA (52), chemogenetic inhibition of the BLA→NAcC pathway did not permanently disrupt the mechanisms underlying the Pavlovian-instrumental interactions involved in the potentiation of cocaine seeking by the conditioned reinforcing properties of cocaine-paired cues. Thus, the effect of chemogenetic inhibition of the BLA→NacC pathway to prevent the impact of cocaine-associated conditioned reinforcement seeking responses was reversible. When Hm4D(GI) rats previously receiving CNO were administered vehicle instead, their seeking responses were potentiated by response-contingent cocaine CS presentation within 6 days of treatment reversal. This observation suggests that the two structures mediate complementary aspects of drug memory (53) and that the BLA→NAcC circuit is necessary for bridging the motivational value of the cocaine-paired cue stored in the BLA with the Pavlovian-instrumental interactive processes supported by the NAcC but is not involved in the learning of either process, as previously suggested (12, 13).

This observation is consistent with the evidence provided here that chemogenetic inhibition of the BLA→NAcC pathway did not influence cocaine seeking per se, since instrumental responding for cocaine under FI15 conditions in the absence of conditioned reinforcement was completely unaffected. This confirms that the acquisition of drug seeking behaviour in anticipation of, and reinforced by, the eventual delivery of a drug infusion (54), does not depend on the BLA or its functional interactions with the NAcC. The observation that cocaine seeking under a FI15 schedule of reinforcement is impervious to the inhibition of the BLA→NAcC pathway at first sight seems inconsistent with the previous demonstration response-contingent optogenetic activation of this pathway in mice supports instrumental responding under continuous reinforcement (36). However, these results taken together with the present data, provide further evidence that instrumental seeking responses are mediated by neural circuits that are dissociable from those mediating directly reinforced taking responses (25), even when emitted within the same behavioural sequence (2).

Finally the present results confirm that the BLA is necessary for the acquisition and early onset performance of cue-controlled cocaine seeking, but not its longer-term maintenance (39). Thus, following extensive training under a second order schedule of reinforcement cue-controlled cocaine seeking was well-established, chemogenetic inhibition of BLA→NAcC pathway, which completely prevented the conditioned reinforcement impact of cocaine cues, had no effect. This observation is in agreement with our previous demonstration that the BLA is necessary for the acquisition of cue-controlled cocaine seeking and the recruitment of dorsolateral striatum-dopamine dependent control over behaviour, but that it is the central amygdala that assumes a critical role in maintaining well established dorsolateral striatum, dopamine-dependent cue-controlled cocaine seeking.

Together the results of the present study reveal the first node of an intricate and shifting amygdalo-striatal circuit that mediates the major influence of pavlovian drug cues acting as conditioned reinforcers to invigorate drug seeking over a protracted time periods in order to obtain intravenous cocaine (38, 48).

## Acknowledgements

DB, MP, BJE and AHM designed the experiment. MP, PB and AHM carried-out the experiments. MP, AHM and DB analysed the data. AHM, MP, BJE and DB wrote the manuscript.

This work, carried at the department of Psychology of the University of Cambridge, was funded by a Programme Grant from the Medical Research Council to BJE and DB (MR/N02530X/1) and a research grant from the Leverhulme Trust to DB (RPG‐2016‐117). AHM was funded by a José Castillejo Grant of the Ministry of Education of Spain and PB was funded by the University of Tours and the University Hospital of Tours, France.

Cocaine hydrochloride and CNO were provided to DB by the NIDA Drug Supply Programme.

