## Supplemental material for "The Basolateral amygdala → Nucleus Accumbens core circuit mediates the conditioned reinforcing effects of cocaine-paired cues on cocaine seeking"

3: CHRU de Tours -  Équipe de Liaison et de Soins en Addictologie & Clinique Psychiatrique Universitaire/ INSERM U1253 iBrain: Imaging & Brain, Tours, France.

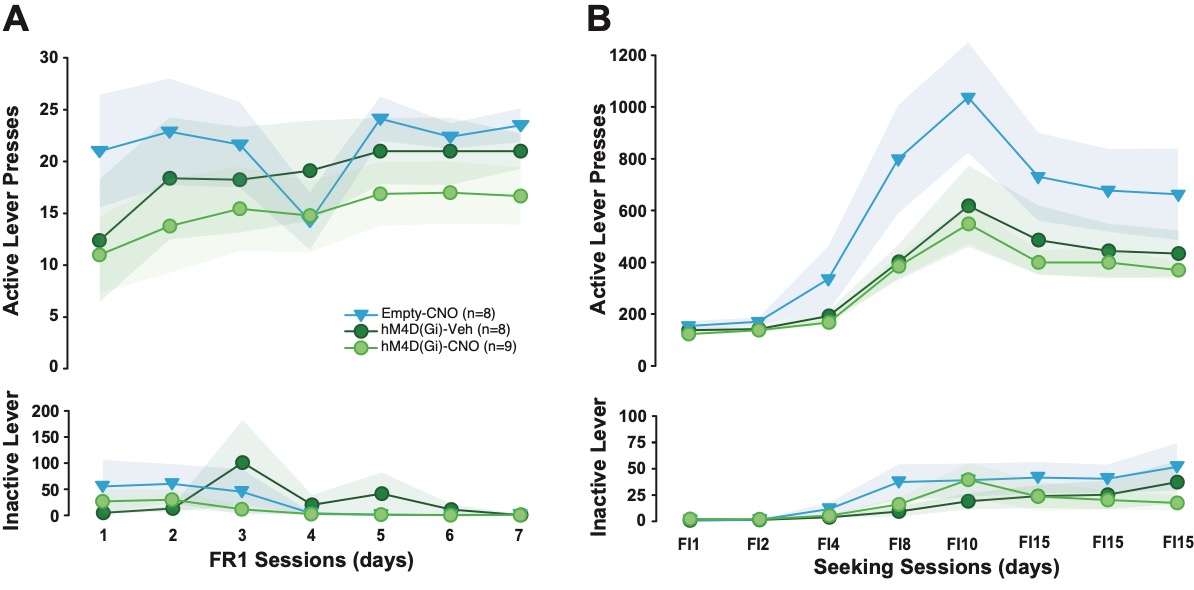

### Supporting Online material

### Figure S1: The cre-mediated expression of an hM4Di inhibitory DREADD or an empty mCherry reporter control virus in the NAcC-projecting BLA neurons had no effect on the acquisition of cocaine self-administration or that of cocaine seeking.

**A)** Active, but not inactive lever presses increased similarly across the three groups over the 7 days of acquisition of cocaine self-administration under fixed ratio 1 (FR1). **B)** Active lever presses increased similarly across groups, from 20 to over 400 over the 8 days of training under fixed interval schedules (FI) of increasing duration, from 1 to 15 minutes.

Empty-CNO, hM4D(Gi)-Veh and Hm4D(Gi)-CNO are shown as blue triangles, dark green and light green circles, respectively.

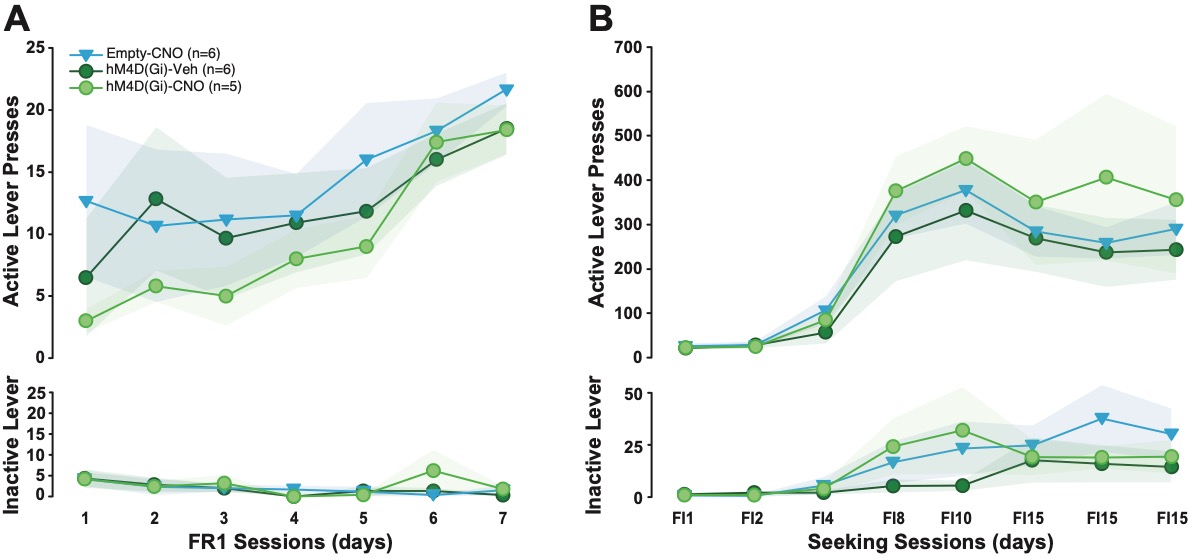

### Figure S2: The expression of an hM4Di inhibitory DREADD or an empty GFP reporter control virus in the anterior insula (AI) had no effect on the acquisition of cocaine self-administration or that of cocaine seeking.

**A)** Active, but not inactive lever presses increased similarly across the three groups over the 7 days of acquisition of cocaine self-administration under fixed ratio 1 (FR1). **B)** Active lever presses increased similarly across groups, from 20 to over 400 over the 8 days of training under fixed interval schedules (FI) of increasing duration, from 1 to 15 minutes.

Empty-CNO, hM4D(Gi)-Veh and Hm4D(Gi)-CNO are shown as blue triangles, dark green and light green circles, respectively.
